# *Runx1* and *Runx2* act in concert to suppress *Wnt/β-catenin*-driven mammary tumourigenesis

**DOI:** 10.1101/2025.03.16.643517

**Authors:** A.I. Riggio, K. Sweeney, R. Shaw, A. Khan, A. Lawlor, N. Ferrari, K. Gilroy, C. Bull, A.L. Young, D. Athineos, H. Hall, F. Ghaffar, C. Nixon, P.D Adams, E.W Roberts, C.J. Miller, P.D. Dunne, K.J. Campbell, E.R. Cameron, K Blyth

**Affiliations:** Cancer Research UK Scotland Institute, Garscube Estate, Switchback Road, Glasgow, G61 1BD, UK; School of Cancer Sciences, University of Glasgow, Garscube Estate, Switchback Road, Glasgow, G61 1QH, UK; School of Biodiversity One Health and Veterinary Medicine, University of Glasgow, Glasgow, G61 1QH, UK; The Patrick G Johnston Centre for Cancer Research, Queen’s University Belfast, Belfast, UK; Sanford Burnham Prebys Medical Discovery Institute, North Torrey Pines Road, La Jolla, CA 92037, USA

**Keywords:** *RUNX1*, *RUNX2*, breast cancer, stemness, tumour initiation, GEMM, WNT, immune

## Abstract

The genes encoding transcription factor *RUNX1* and its binding partner *CBFB* have been reported to be mutated in human breast cancer. Here, we provide evidence that *Runx1* loss of function results in accelerated disease onset and tumour development in mouse models of breast cancer, in keeping with a tumour suppressor role for RUNX1 in this disease setting. Combined deletion of *Runx1* and the related family member *Runx2* resulted in mammary epithelial cells becoming exquisitely sensitive to WNT-driven transformation, with the emergence of multiple tumours early in life. Clonogenic assays indicated that *Runx1* ablation induced a stem cell like phenotype in mammary epithelial cells, whilst transcriptome analysis demonstrated activation of multiple oncogenic pathways, especially when *Runx2* was co-deleted. Interestingly, altered *Runx* expression in the mammary epithelium also drove profound alterations in the tumour microenvironment, impacting the immune landscape. These results highlight that *Runx1* restricts some forms of breast cancer and inhibits the full oncogenic potential of aberrant WNT signalling. Loss of *Runx2* itself did not result in tumour promotion, yet the dramatic effects of combined *Runx1* and *Runx2* loss suggest that *Runx2* can substitute for *Runx1* in dampening the oncogenic effects of WNT signalling.

## INTRODUCTION

Breast cancer is the commonest malignant disease affecting women and, despite significant improvement in survival rates, remains a major cause of mortality (1). Rather than a single disease, breast cancer is a heterogeneous group of tumours defined by anatomical origin and classified by histopathological, clinical and molecular features (2). Based on receptor status, breast tumours are categorised into three subtypes: hormone receptor positive (oestrogen receptor, ER; progesterone receptor, PR), human epidermal growth factor receptor type 2 (HER2) positive and triple negative (TN, ER-/PR-/HER2-) (3). Molecular profiling has helped stratify these distinct diseases into separate intrinsic subtypes that largely, but not entirely, map to the well-established disease categories (4–6). While clinically useful, it is oversimplistic to assume each of the histological subtype is always defined by a common genetic landscape and indeed, molecular heterogenicity exists within well characterised types of breast cancer.

The *RUNX* genes (*RUNX1, −2* and *−3*) encode transcription factors that bind to a common co-factor, CBFβ, and modulate expression of many target genes, either directly through binding to the RUNX DNA consensus sequence or indirectly through protein-protein interactions with other partners (7). The emergence of these genes as players in breast cancer is evident (8), yet their functionality in tumour development and progression is complex and contradictory (9, 10). This is particularly true for *RUNX1* where changes in gene expression have unpredictable outcomes on cell survival, differentiation and proliferation. Large-scale mutational and expression studies indicate that loss of *RUNX1/CBFB* is associated with a significant proportion of breast cancers. *RUNX1* is a frequent mutational target, with genetic changes predicted to interfere with its DNA binding activity or binding to CBFβ, resulting in reduced function (11–13). *CBFB* is also frequently mutated, supporting the notion that RUNX1 acts as a tumour suppressor in breast cancer (11, 12). These putative loss-of-function mutations in *RUNX1* and *CBFB* were overrepresented in luminal breast cancers, suggesting that the RUNX1/CBFβ complex may restrict some subtypes (7). Mutational studies are supported by mRNA expression profiling and protein levels, where reduced *RUNX1* is associated with breast cancer metastasis and high-grade tumours (14, 15). We, and others, have previously demonstrated that *Runx2* (but not *Runx3*) is expressed in the mammary epithelium, although at lower levels than *Runx1* (16, 17). Importantly, loss-of-function mutations of *CBFB* would ablate both RUNX1 and RUNX2 activity. Although the role of *RUNX2* in breast cancer has not been fully explored, we showed that *Runx2* supports mammary reconstitution and stem cell function (18). In contrast to its tumour suppressor role, transcriptome analyses showed *RUNX1* upregulation in some subtypes (19, 20) and enhanced protein expression is associated with poor outcome in TN tumours (21). Therefore, with an obvious, yet unresolved role in breast cancer, investigation of *Runx* genes in physiologically relevant models is warranted.

As genetically engineered mouse models (GEMMs) have proved useful in dissecting the interaction of oncogenic pathways (22), we sought to assess the putative tumour suppressor functions of *Runx1* using *in vivo* models. Loss of *Runx1* accelerated disease in specific oncogenic backgrounds including a model with activated WNT signalling, an important alteration in breast cancer. Components of the WNT pathway, including β-catenin, are frequently overexpressed across breast cancer subtypes, associated with a stem-like phenotype and poorer outcomes (23–25). We show functional overlap between *Runx1* and *Runx2* in restricting *Wnt/β-catenin*-driven tumorigenesis, with *Runx*-deficient β-catenin-expressing mammary cells being exquisitely prone to tumour formation. *Runx* loss increases stemness of mammary cells and unleashes the oncogenic potential of deregulated WNT/β-catenin signalling. Furthermore, loss of *Runx1* in the epithelium alters the tumour microenvironment.

## RESULTS

### *Runx1* loss facilitates early tumour onset in mouse models of mammary cancer

To investigate the possible tumour suppressor role of RUNX1 in breast cancer, we deleted *Runx1* in preclinical mouse models. *MMTV-PyMT* is a well characterised breast cancer model (26), transcriptionally aligning with the luminal B subtype (27) and exhibiting early mammary hyperplasia progressing to highly aggressive tumours. To explore *Runx1* loss on *PyMT*-driven tumourigenesis, we generated *MMTV-PyMT;MMTV-Cre*;*Runx1^fl/fl^*(hereafter *PyMT+/R1KO*) mice and *MMTV-PyMT*;*MMTV-Cre;Runx1^wt/wt^*(hereafter *PyMT+/WT*) controls. Given that *PyMT*-driven tumorigenesis and *Cre*-mediated recombination in this model could occur asynchronously and independently, a conditional *tdRFP* reporter (28) was introduced as a surrogate for *MMTV-Cre* mediated *Runx1* allele excision (Supplement Fig. 1A). Loss of *Runx1* significantly (*P* = 0.0001) accelerated formation of mammary tumours in *PyMT+/R1KO* mice (average time to tumour onset of 50 days, ± 10.9) compared to *PyMT+/WT* mice (average time to tumour onset of 61 days, ± 11.5), indicating that RUNX1 normally restricts the oncogenic pathways activated by *PyMT* (Fig. 1A). No significant difference in tumour volume between cohorts was observed at tumour onset (Supplement Fig. 1A). To explore if *Runx1* deletion was manifesting a selective advantage to the epithelial population primed for transformation, we assessed the proportion of *tdRFP*-positive cells in MMECs isolated from *PyMT+/WT* and *PyMT+/R1KO* mice at tumour notice, as well as from *PyMT-*negative counterparts at ∼60 days. Our results showed a significantly (*P* < 0.01) increased percentage of *tdRFP* positivity in *Runx1*-deleted MMECs, regardless of the status of the *PyMT* oncogene (Fig. 1B).

**Fig. 1.**
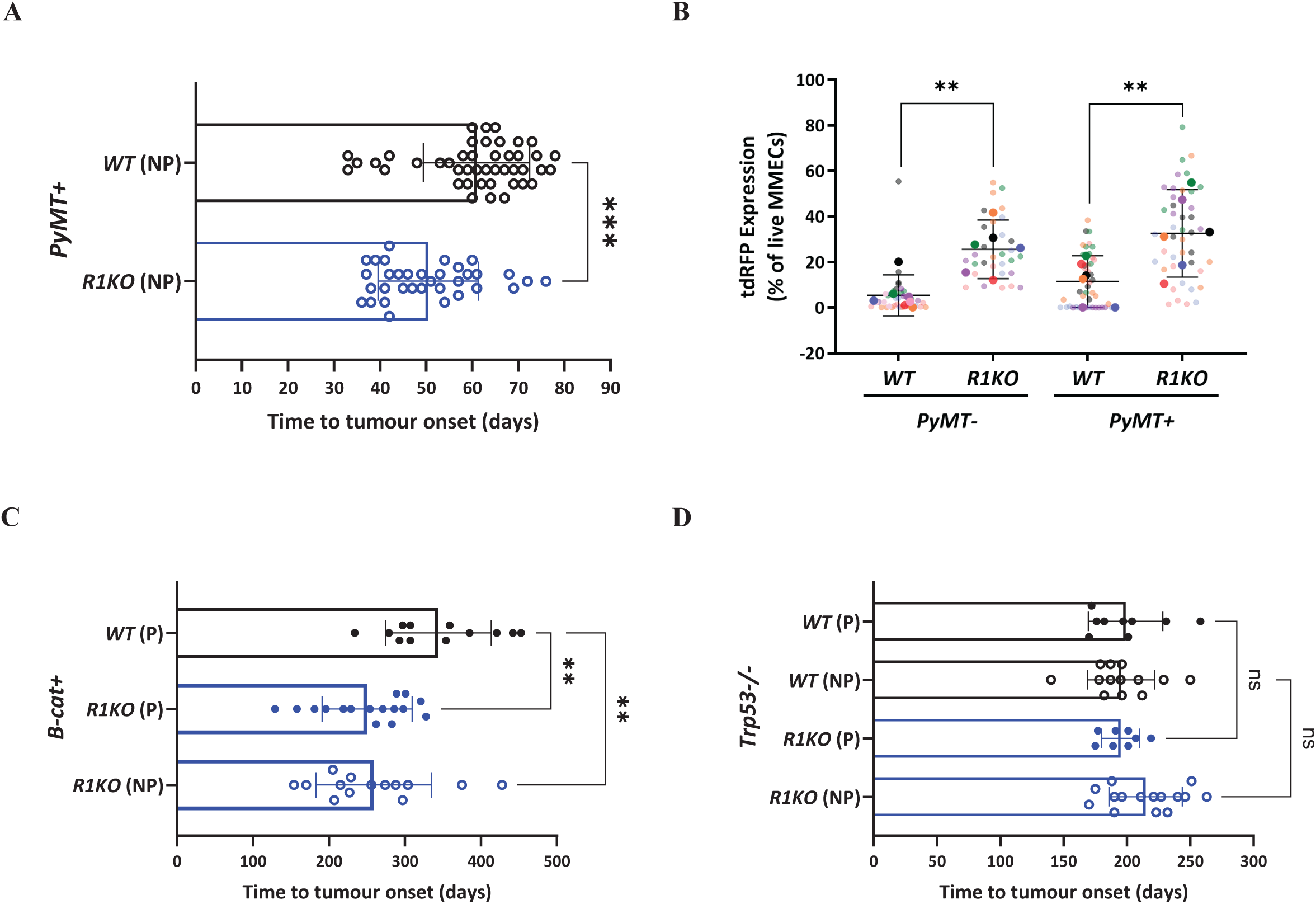
Loss of *Runx1* facilitates early tumour onset in preclinical models of breast cancer. A. Time to tumour onset of *PyMT+* mice wild type (*n* = 44; *WT*) or deleted for *Runx1* (*n* = 34; *R1KO*). Average time to tumour onset with standard deviation shown: 61 days (± 11.5) for *WT;* 50 days (± 10.9) for *R1KO* mice. Each data point represents an individual mouse. NP: nulliparous. Statistical analysis performed with two-tailed unpaired t-test, \*\*\**P* = 0.0001. B. Flow cytometric analysis of *tdRFP* expression in MMECs from *PyMT+/WT* (*n* = 6) and *PyMT+/R1KO* (*n* = 6) mice at tumour notice, and *PyMT*-negative controls (*PyMT-/WT, n* = 7; *PyMT-/R1KO, n* = 6) at ∼60 days. Each data point represents percentage of *tdRFP*-positive MMECs, with smaller points representing individual glands (5-7 glands per mouse, see methods) and larger points representing the means from each mouse. Glands isolated from the same mouse are indicated by the same colour. Statistical analysis performed on the mean percentage of *tdRFP*-positive MMECs with ordinary one-way ANOVA with Sidak’s multiple comparisons test, \*\**P* < 0.01. C. Time to tumour onset of *BLG-Cre;Ctnnb1^wt/lox(ex3)^* (*B-cat+*) mice wild type (*n* = 12; *WT*) or deleted for *Runx1* (*n* = 16; parous *R1KO* and *n* = 14; nulliparous *R1KO*). Average time to tumour onset with standard deviation shown: 344 days (± 69.6) for parous *WT*; 250 days (± 59.3) for parous *R1KO*; 259 days (± 75.9) for nulliparous *R1KO* mice. Each data point represents an individual mouse. Parous (P) mice, filled circles; nulliparous (NP) mice, open circles. Statistical analysis performed with ordinary one-way ANOVA test with Dunnett’s multiple comparisons test, \*\**P* < 0.01. D. Time to tumour onset in *BLG-Cre;Trp53^fl/fl^* (*Trp53-/-*) parous (P) and nulliparous (NP) mice wild type (*n* = 9 and *n* = 13, respectively; *WT*) and deleted for *Runx1* (*n* = 8 and *n* = 15, respectively; *R1KO*). Each data point represents an individual mouse. Average time to tumour onset with standard deviation shown. *WT*-P: 199 days (± 29.4); *WT*-NP: 195 days (± 26.7); *R1KO-*P: 195 days (± 15); *R1KO*-NP: 215 days (± 28.9). Statistical analysis performed with ordinary one-way ANOVA test with Sidak’s multiple comparisons test; ns, non-significant (*P* > 0.05).

We next tested the effects of *Runx1* deletion in a novel model of *Wnt/β-catenin*-driven breast cancer by crossing mice carrying a stabilised form of β-catenin (*Ctnnb1^wt/lox(ex3)^*) (29) onto the *BLG-Cre* line (30). *BLG-Cre;Ctnnb1^wt/lox(ex3)^* mice (hereafter *B-cat+*), either *Runx1*- proficient (hereafter *B-cat+/WT*) or *Runx1*-deficient (hereafter *B-cat+/R1KO*), were generated (Supplement Fig. 1B). Loss of *Runx1* resulted in a significant (*P* < 0.01) acceleration of *Wnt/β-catenin*-driven tumorigenesis, with *B-cat+/R1KO* mice developing tumours at an average onset of 250 days (± 59.3) as opposed to the average 344 days (± 69.6) of *B-cat+/WT* animals (Fig. 1C). Of note, *Runx1* deletion removed the need for parity required to maximise BLG-Cre expression, with 16 out of 20 (80%) nulliparous *B-cat+/R1KO* animals aged to 325 days presenting with palpable tumours compared to only 1 out of 12 (8%) nulliparous *B-cat+/WT* mice aged to the same time. Furthermore, tumour onset in nulliparous *B-cat+/R1KO* animals occurred at the same latency as the corresponding parous cohort (259 days, ± 75.9; Fig. 1C). There was no difference in tumour volume between cohorts at tumour notice (Supplement Fig. 1B). Thus, *BLG-Cre*-mediated loss of *Runx1*, in combination with constitutive activation of WNT/β-catenin, significantly accelerates mammary tumourigenesis.

Interestingly, despite the tumour promoting effects of *Runx1* deletion in both *MMTV-PyMT* and *Wnt/β-catenin* models, *Runx1* loss did not accelerate tumour development in a *BLG-Cre;Trp53^fl/fl^* GEMM (Fig. 1D & Supplement Fig. 1C), pointing towards possible redundancy in the tumour suppressive actions of *Trp53* and *Runx1*.

### Genetic loss of *Runx1* and *Runx2* dramatically accelerates *Wnt/β-catenin*-driven mammary tumour development

Given the clinical relevance of WNT/β-catenin in breast cancer (31, 32) and the co-occurrence of *WNT/RUNX* gene alterations (33), we further investigated the tumour suppressor phenotype shown by *Runx1* loss in the *Wnt/β-catenin*-driven model. It was previously reported that RUNX1 has a role in the differentiation of mature ER+ luminal mammary epithelial cells (34), whilst our group showed that RUNX2 is associated with mammary stemness (18). Given the auto and cross regulation between these proteins and that through a common binding site they regulate the same targets, we sought to understand the effects of combined *Runx1* and *Runx2* loss, hypothesising that *Runx2* might substitute for *Runx1* loss in restricting the oncogenic potential of WNT/β-catenin. Accordingly, crossing *B-cat+* mice with *Runx1^fl/fl^*;*Runx2^fl/fl^*animals (hereafter *B-cat+/DKO*) resulted in a dramatic acceleration of disease onset (*P* < 0.0001), with nulliparous *B-cat+/DKO* females developing lesions with a median tumour-free age of 56 days (± 14.9) compared to both parous *B-cat+/R1KO* (median tumour-free age of 266 days ± 59.3) and *B-cat+/WT* (median tumour-free age of 330 days ± 69.6) mice (Fig. 2A and Supplement Fig. 1D). Indeed, tumour development was too rapid to permit even one round of pregnancy in *B-cat+/DKO* females. Loss of *Runx2* alone (*B-cat+/R2KO*) did not affect the rate of tumour onset compared to *B-cat+/WT* mice (Fig. 2A), indicating that the extreme phenotype was only observed with loss of both genes.

**Fig. 2.**
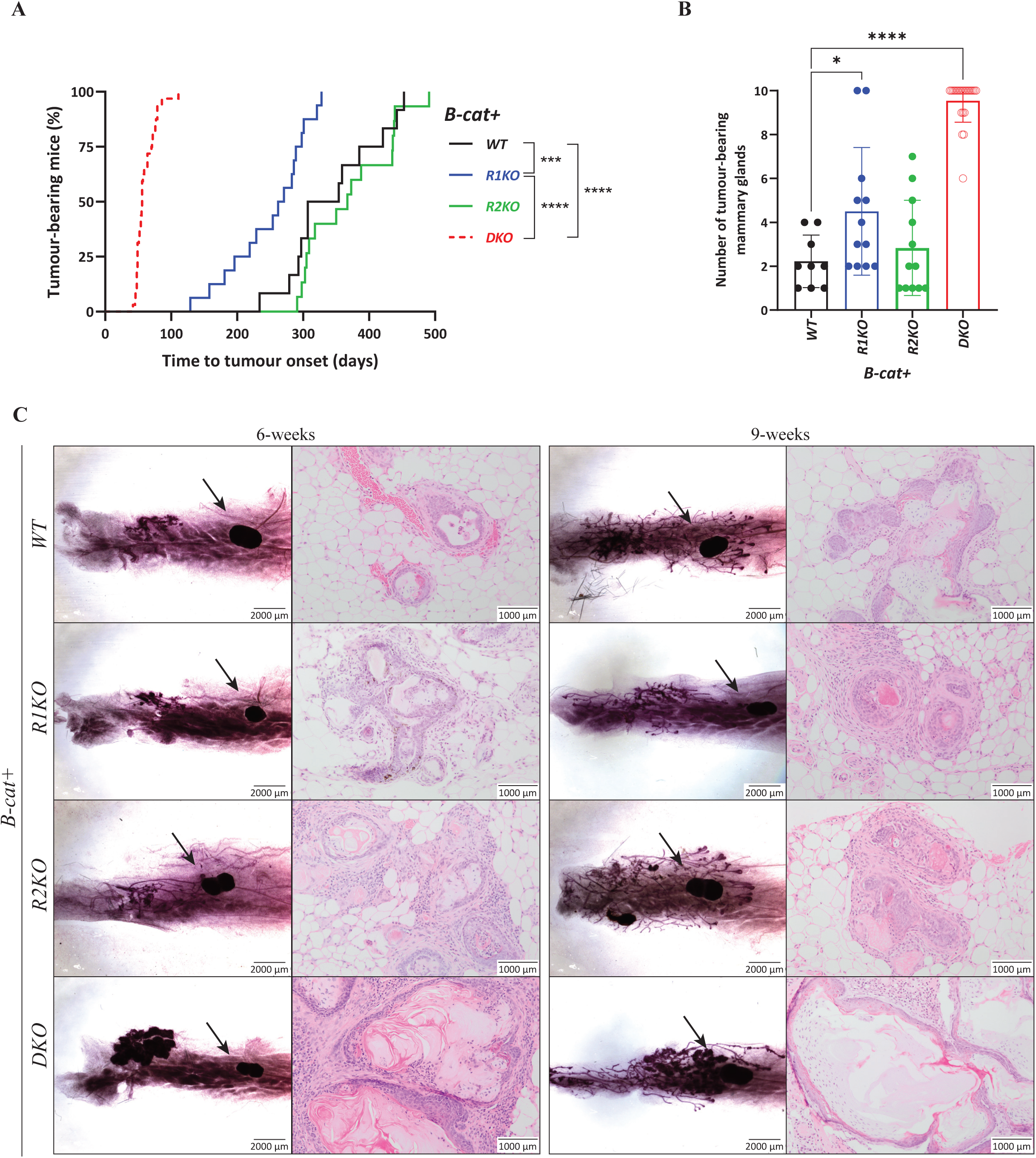
Combined loss of *Runx1* and *Runx2* dramatically accelerates *Wnt/*β*-catenin*- driven mammary tumour development. A. Time to tumour onset in *B-cat+* mice wild-type for *Runx* (*n* = 12; *WT* as in Fig. 1C), *Runx1^fl/fl^* (*n* = 16; *R1KO* as in Fig. 1C), *Runx2^fl/fl^* (*n* = 15; *R2KO*) and *Runx1^fl/fl^;Runx2^fl/fl^*(*n* = 32; *DKO*). Parous (P) mice, solid lines; nulliparous (NP) mice, dotted lines. Median survival of tumour-free mice with standard deviation shown: *WT*-P mice, 330 days (± 69.6); *R1KO*-P mice, 266 days (± 59.3); *R2KO*-P mice, 367 days (± 65.4); *DKO*-NP mice, 56 days (± 14.9). Statistical analysis performed with log-rank (Mantel-Cox) test, \*\*\**P* < 0.001; **** *P* < 0.0001. B. Number of tumour-bearing mammary glands in *B-cat+* cohorts from Fig. 2A harvested at clinical endpoint. Each data point represents an individual mouse. Parous (P) mice, filled circles; nulliparous (NP) mice, open circles. *WT*-P (*n* = 9); *R1KO*-P (*n* = 12); *R2KO*-P (*n* = 12); *DKO*-NP (*n* = 24). Statistical analysis performed with ordinary one-way ANOVA with Dunnett’s multiple comparisons test, \**P* < 0.05; \*\*\*\**P* < 0.0001. C. Wholemounts and H&E images of abdominal mammary glands from 6- and 9-weeks old *B-cat+* cohorts. One representative wholemount and H&E image is shown per genotype (of *n* = 5-7 mice per genotype). Scale bars of wholemounts = 2mm; H&E images = 1mm. Arrow illustrates lymph node.

Examination of *B-cat+/DKO* mice showed a very distinct presentation, with multiple, sometimes coalescing tumours, as opposed to *B-cat+/WT, B-cat+/R1KO* and *B-cat+/R2KO* animals presenting with a smaller number of larger lesions (Supplement Fig. 2A). Accordingly, a significant increase in the number of tumour bearing glands was observed in *B-cat+/DKO* (*n* = 9/10 on average; *P* < 0.0001) compared to *B-cat+/WT* (*n* = 2/10 on average) mice (Fig. 2B). A modest, but significant, difference was also observed between *B-cat+/R1KO* (*n* = 4/10 on average; *P* < 0.05) and *B-cat+/WT* controls (Fig. 2B). This pathological profile, combined with faster onset of disease, indicates that the threshold for *Wnt/β-catenin-*driven mammary tumour development is significantly lower with simultaneous loss of *Runx1* and *Runx2* activity.

To track early neoplastic changes, we examined wholemounts in virgin females. Activation of *Wnt/β-catenin* signalling resulted in mild hyperplasia in control *B-cat+/WT* mice, as well as in *B-cat+/R1KO* and *B-cat+/R2KO* cohorts (Fig. 2C). However, *B-cat+/DKO* mice exhibited dramatic changes, with hyperplastic outgrowths and disrupted architecture as confirmed by wholemounts and H&E images (Fig. 2C). As this phenotype was not observed in the absence of activated *β-catenin* (Supplement Fig. 2B), loss of RUNX function alone does not overtly interfere with mammary gland development. Rather, RUNX proteins act in concert to suppress *Wnt/β-catenin*-driven expansion of a mammary epithelial cells.

### Loss of RUNX expands a population of mammary epithelial cells with stem-like properties in the presence of activated WNT/β-catenin signalling

Given the early changes described above and the increased tumour-bearing glands at endpoint in *B-cat+/DKO* animals (Fig. 2B), we speculated that combined loss of *Runx1* and *Runx2* could alter stem/progenitor population(s) to favour *Wnt/β-catenin*-driven preneoplastic expansion and outgrowth of mammary tumours. Thus, we isolated MMECs from 7-9 weeks old *B-cat+* mice and performed colony formation cell assay. Stingl *et al.* (35) demonstrated that fractions enriched for cells capable of mammary gland regeneration *in vivo* (putative stem cells referred to as mammary repopulating units, MRUs) can be distinguished from the more numerous cells capable of colony formation *in vitro* (putative progenitor cells referred to as mammary colony forming units, Ma-CFU) by the type of colonies formed. While Ma-CFU give rise to spherical acinar structures, MRUs generate solid, irregular-shaped colonies in Matrigel culture. Interestingly, *B-cat+/DKO* and *B-cat+/R1KO* cells showed significantly reduced (*P* < 0.0001) acinar-like colonies in favour of a significant increase (*P* < 0.0001) in solid-like colonies compared to *B-cat+/WT* MMECs (Fig. 3A). These results demonstrate that loss of RUNX function expands a pool of cells enriched for stem-like properties.

**Fig. 3.**
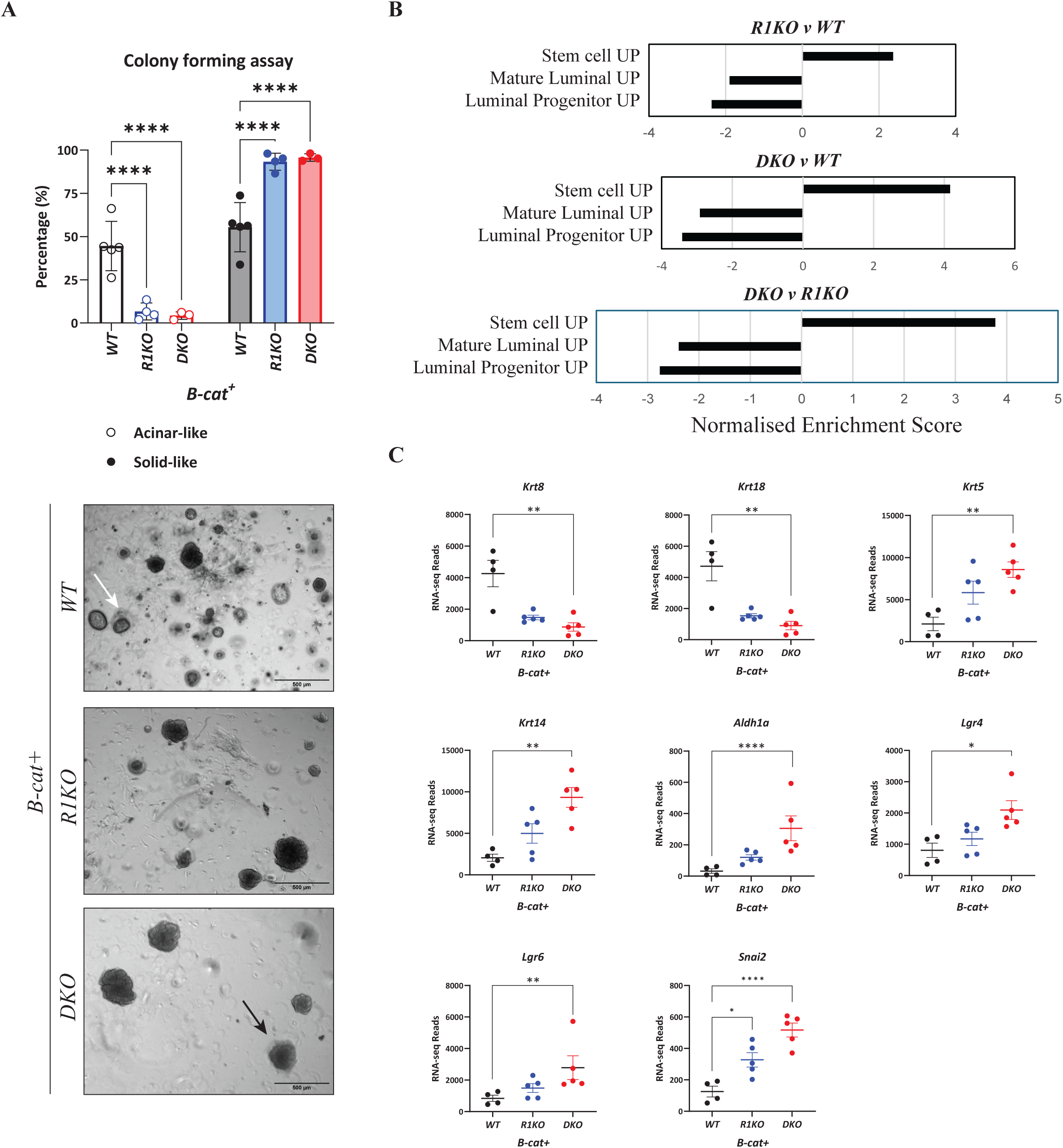
Loss of RUNX function alters mammary stem/progenitor populations. A. Graph (top) and representative pictures (bottom) of colony-forming cell assay showing relative proportions of acinar- and solid-like colonies from MMECs isolated from 7-9 weeks old *B-cat^+^*cohorts; *Runx1* wild type (*n* = 5; *WT*); *R1KO* (*n* = 4); *DKO* (*n* = 3). Mean with standard error of *n* ≥ 3 independent experiments, each with 12 technical replicates per condition. One representative image shown per cohort (*n* ≥ 3 biological replicates per cohort). Acinar and solid-like colonies depicted by white and black arrows, respectively. Statistical analysis performed with two-way ANOVA with Dunnett’s multiple comparisons test, \*\*\*\**P* < 0.0001. Scale bar, 500µm. B. Analysis of mammary RNA-seq signatures derived from gene set enrichment analysis (GSEA) of FACS-sorted *tdRFP-*positive MMECs from 9-weeks old *B-cat^+^* females comparing three conditions: *Runx1* knockout (*R1KO; n* = 5) versus wildtype for *Runx* genes (*WT; n* = 4), *Runx1/Runx2* knockout (*DKO; n* = 5) versus *WT*, and *DKO* versus *R1KO*. The gene sets analysed include LIM Mammary Stem Cell Up (https://www.gsea/msigdb.org/gsea-msigdb/mouse/geneset/LIM_MAMMARY_STEM_CELL_UP.html), LIM Mammary Mature Luminal Up (https://www.gsea-msigdb.org/gsea/msigdb/mouse/geneset/LIM_MAMMARY_STEM_CELL_UP.html), and LIM Mammary Luminal Progenitor Up (https://www.gsea-msigdb.org/gsea/msigdb/mouse/geneset/LIM_MAMMARY_STEM_CELL_UP.html). Each bar represents the normalised enrichment score (NES) for a specific gene set under each comparison, with higher NES values indicating stronger enrichment of the gene set in the corresponding condition. C. Normalised gene expression levels of selected genes in FACS-sorted *tdRFP-*positive MMECs as described for Fig. 3B. Individual data points represent expression levels per sample, with central line indicating the median. Gene expression normalised for sequencing depth, with significant differences between conditions assessed with Wald test and shown as p-values adjusted for multiple testing with Benjamini-Hochberg method. Both procedures performed with DESeq2; \**P* < 0.05; \*\**P* < 0.01; \*\*\**P* < 0.001; \*\*\*\**P* < 0.0001.

To help identify MMECs with simultaneous *Wnt/β-catenin* activation and *Runx* loss, *B-cat+* mice (*WT*, *R1KO* and *DKO*) were crossed with the *tdRFP* line. RNA-seq analysis was then performed on flow sorted *tdRFP-*positive MMECs lacking *Runx1*, or both *Runx1* and *Runx2*. Interestingly, an enrichment of gene sets aligning with mammary stem cells and negatively associated with mature luminal and progenitor populations, was observed in MMECs isolated from *B-c*at+/*R1KO* mice compared to *B-cat*+/*WT* animals (Fig. 3B). However, the same expression profile showed a stronger enrichment in MMECs from *B- cat+/DKO* mice (Fig. 3B), indicating that combined loss of *Runx1* and *Runx2* further shifts MMECs to a stem-like state. Accordingly, expression levels of luminal markers (i.e. *Krt8* and *Krt18*) were reduced and basal markers (i.e. *Krt5* and *Krt14*) increased in *B-cat+/DKO* cells compared to their *WT* counterpart (Fig. 3C), in line with a shift to more solid-like colonies and the gene expression signature of MRUs (35). In addition, genes important in cancer stem cells, epithelial-to-mesenchymal transition and aggressiveness, such as *Aldh1a*, *Lgr4*, *Lgr6* and *Snai2* (36–39), were mildly and markedly increased in *R1KO* and *DKO* cells, respectively, compared to *B-cat+/WT* MMECs (Fig. 3C). Altogether, these data indicate a shift towards an earlier lineage stage when *Wnt/β-catenin* activation occurs with simultaneous *Runx* loss.

### *Runx* gene loss increases mammosphere formation in murine epithelial cells

To study RUNX function in a more tractable system, we used CRISPR-Cas9 to target *Runx1* and *Runx2* separately and in combination, in the non-tumorigenic, mammary HC11 cell line (40) (Supplement Fig. 3A-C). There was no obvious change in phenotype or growth characteristics when cells with *Runx1* and/or *Runx2* loss were grown in 2D conditions (Supplement Fig. 3D), nor when *Runx1* was overexpressed (Supplement Fig. 3B-D). However, *Runx1* loss significantly increased mammosphere formation when HC11 cells were grown in suspension (Fig. 4A), whilst the reverse was observed upon *Runx1* overexpression (Fig. 4B). By contrast, loss of *Runx2* reduced mammosphere number (Fig. 4C), in line with previous results showing *Runx2* driving a stem cell phenotype (18). Combined loss of *Runx1* and *Runx2* increased mammosphere formation similar to that with *Runx1* loss (Fig. 4C), emphasising that RUNX1 is the predominant factor moving cells out of the stem cell compartment. Given the exquisite relationship between *Runx* and *Wnt/β-catenin* signalling revealed above, recombinant WNT3A ligand was added to mammosphere cultures. While WNT3A increased mammosphere formation in HC11 cells wild-type for *Runx*, as expected, it further potentiated stemness of *R1KO* and *DKO* clones (Fig. 4D). Altogether, these results suggest complementary effects of *Wnt/β-catenin* signalling and loss of *Runx1* (and *Runx2*) on mammosphere formation.

**Fig. 4.**
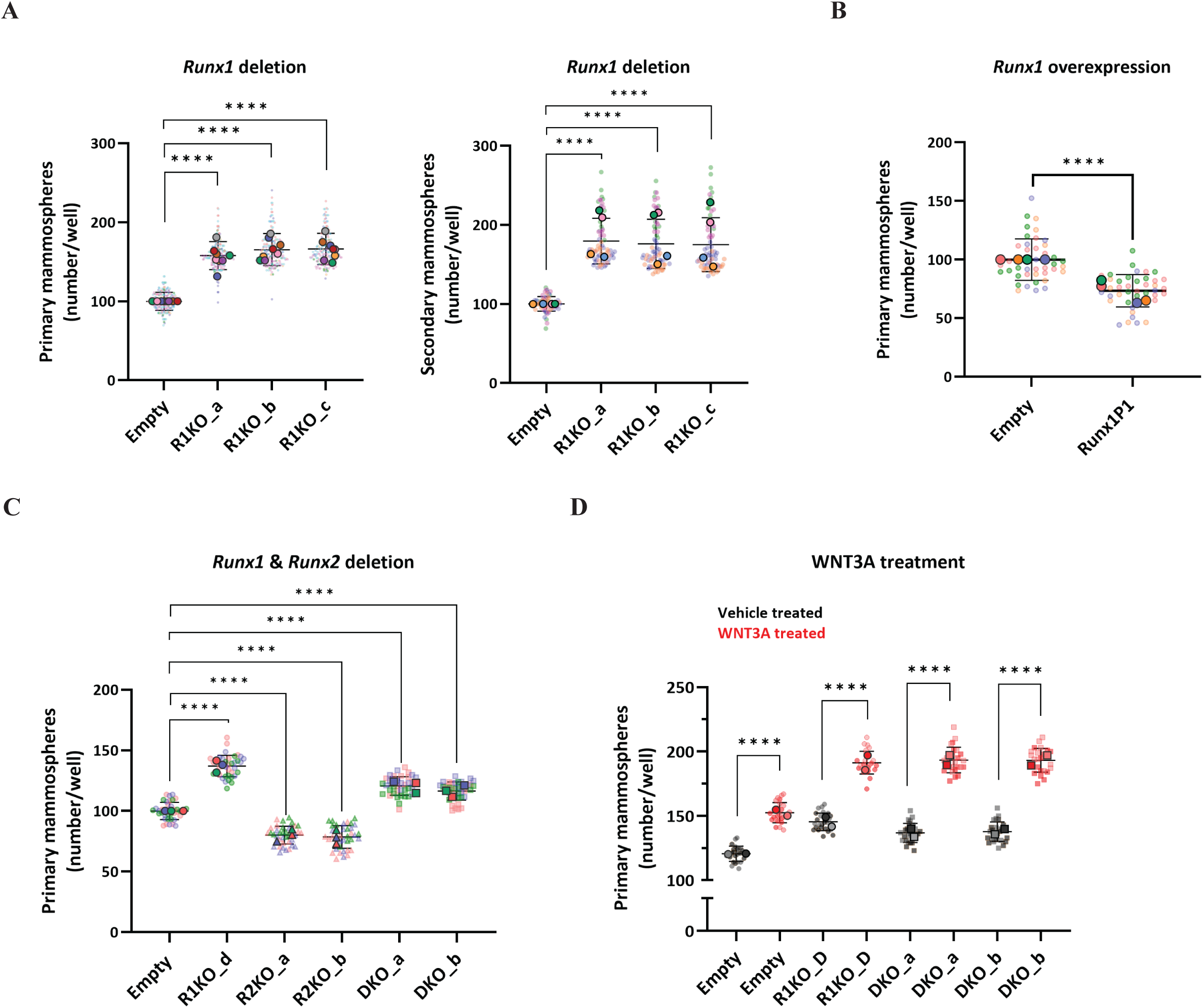
*Runx* gene loss increases mammosphere formation in HC11 cells. A. Scatter dot-plot of primary (left) and secondary (right) mammospheres of HC11 cells with empty vector (Empty) or independent CRISPR/Cas9-deleted *Runx1* clones (R1KO_a, R1KO_b, R1KO_c). Results show normalised mammosphere counts per well relative to the average mammosphere count in control (Empty) group. Smaller points represent technical replicates, larger points the mean of each experimental repeat with coloured points used to differentiate experimental replicates (also for B-D below). For primary mammospheres, there were 12-48 technical replicates per condition per experimental replicate (*n* = 8). Error bars indicate normalised mean of all technical replicates ± standard deviation (Empty, 100 ± 11.5; *R1KO_a*, 158.1 ± 17.9; *R1KO_b*, 165.4 ± 20.7; *R1KO_c*, 166.3 ± 20.0). For secondary mammospheres, there were 12-24 technical replicates per condition for each experimental repeat (*n* = 4). Error bars indicate normalised mean of all technical replicates ± standard deviation (Empty, 100.1 ± 9.6; *R1KO_a*, 179.1 ± 28.9; *R1KO_b,* 175.6 ± 31.1; *R1KO_c*, 174.5 ± 34.1). Statistical analysis performed with ordinary one-way ANOVA with Dunnett’s multiple comparisons test, \*\*\*\**P* < 0.0001. B. Scatter dot-plot of primary mammospheres of HC11 cells overexpressing *Runx1* (*Runx1P1*) or empty vector control (Empty). Results show mammosphere counts relative to the average mammosphere count in control group (Empty). Error bars indicate normalised mean of all technical replicates ± standard deviation (Empty, 100 ± 18.6; *Runx1P1*, 73.5 ±14.3) and data are representative of *n* = 4 experimental repeats. For each experiment, there were 8-12 technical replicates per condition. Statistical analysis performed with two-tailed unpaired t-test, \*\*\*\**P* < 0.0001. C. Scatter dot-plot of primary mammospheres of HC11 cells first deleted for *Runx2* and then transduced with lentiCRISPRv2-Neo empty vector (Empty) or containing a *Runx1*-targeting gRNA plasmid to generate an independent *Runx1*-deleted (*R1KO_d*), two *Runx2*-deleted (*R2KO_a*, *R2KO_b*) and two *Runx1/Runx2*-deleted (*DKO_a, DKO_b*) clones. Results show normalised mammosphere counts per well relative to the average mammosphere count in control group (empty). Error bars are normalised mean of all technical replicates with standard deviation (Empty, 100 ± 7.4*; R1KO_d,* 137.1 ± 9.1; *R2KO_a*, 80.1 ± 7.6; *R2KO_b*, 78.6 ± 9.6; *DKO_a,* 120.6 ± 7.9; *DKO_b,* 116.4 ± 7.7) and data are representative of *n* = 3 experimental repeats. For each experiment, there were 12 technical replicates per condition. Statistical analysis performed with ordinary one-way ANOVA with Dunnett’s multiple comparisons test, \*\*\*\**P* < 0.0001. D. Scatter dot-plot of primary mammospheres of HC11 cells as for C in the presence or absence of recombinant WNT3A ligand. Error bars show mean of all technical replicates ± standard deviation for vehicle treated (Empty, 120.5 ± 6.2; *R1KO_d*, 145.4 ± 7.0; *DKO_a*, 136.8 ± 7.7; *DKO_b*, 137.7 ± 7.9) and WNT3A treated (Empty, 152.5 ± 8.1; *R1KO_d*, 191.3 ± 9.0; *DKO_a*, 193.4 ± 10.2; *DKO_b,* 193.1 ± 9.5) clones, of *n* = 2 experimental repeats. For each experiment, 12 technical replicates were tested per condition. Statistical analysis performed with ordinary one-way ANOVA with Sidak’s multiple comparisons test, \*\*\*\**P* < 0.0001.

### Loss of *Runx1* and *Runx2* results in the activation of oncogenic pathways

To understand the pathways driving accelerated *Wnt/β-catenin*-driven oncogenic development in the absence of both *Runx* genes, we explored RNA-seq data of the *tdRFP-*positive sorted MMECs. In line with the milder phenotype and modest reduction in tumour development observed between *B-cat+/WT* and *B-cat+*/*R1KO* groups, only 88 differentially expressed genes reached significance upon *Runx1* loss. By contrast, 1312 genes were significantly different between *B-cat+/WT* and *B-cat+*/*DKO* cohorts, demonstrating a greatly altered gene expression in the absence of both *Runx* genes (Fig. 5A). Of note, 75/88 (85%) significantly altered genes within the *B-cat+*/*R1KO* set were shared with the *B-cat+/DKO* cohort (Fig. 5A). When we interrogated gene set enrichment analysis and the Hallmark data sets using a significance cut- off of P < 0.01 and a false discovery rate of P < 0.05, 41 Hallmark gene sets were found to be enriched in the *B-cat+/DKO* dataset compared to *B-cat+/WT,* whilst only 28 were significantly enriched in the *B-cat+/R1KO* group relative to the *B-cat+/WT*. Notably, the majority of these gene sets (24/28) were also shared with the *B-cat+/DKO* cohort (Fig. 5A).

**Fig. 5.**
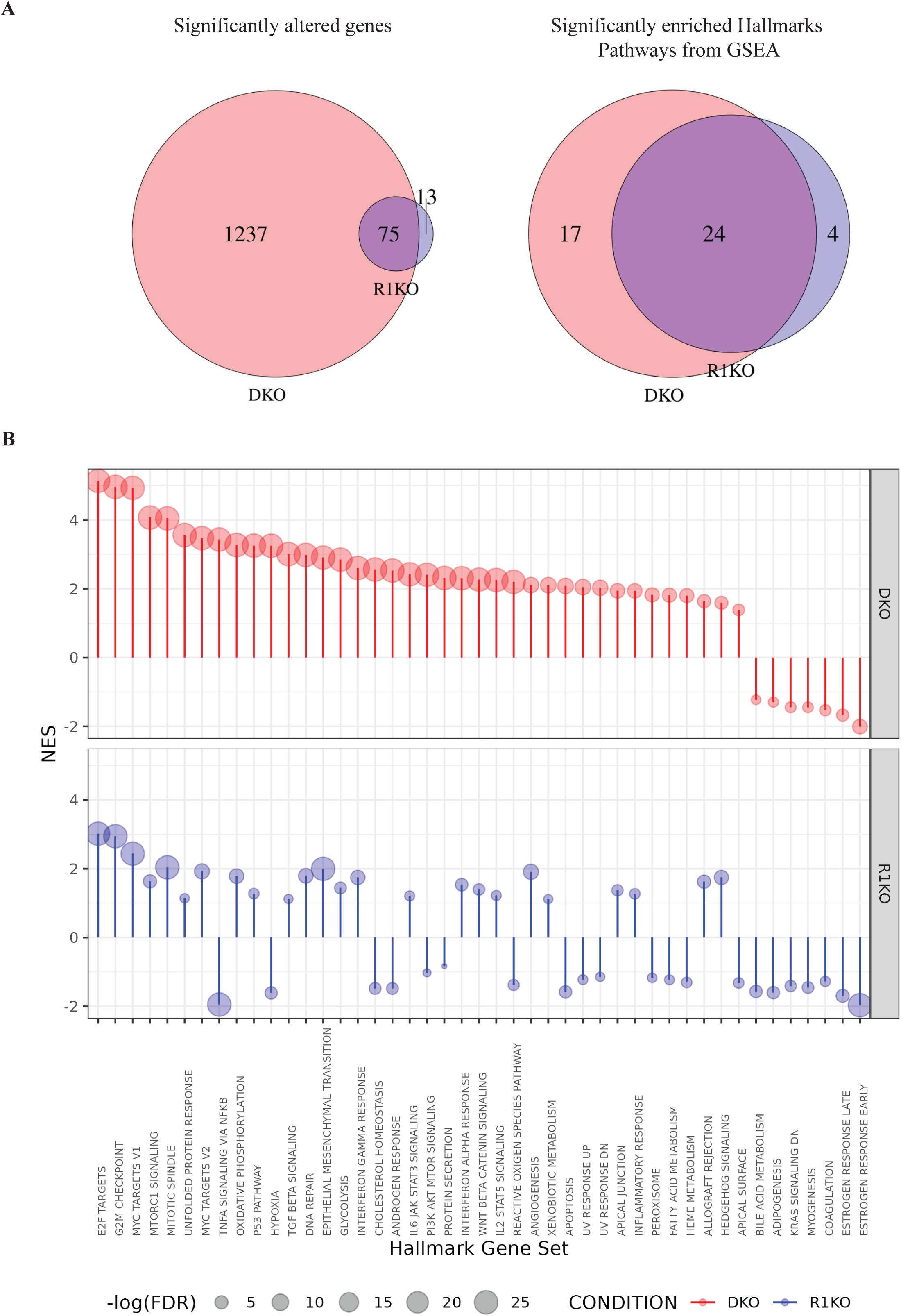
Combined loss of *Runx1* and *Runx2* results in the activation of key oncogenic pathways. RNA-seq analysis of MMECs isolated from 9-weeks old *B-cat^+^/WT* (*n* = 4), *B-cat^+^/R1KO* (*n*= 5) and *B-cat^+^/DKO* (*n* = 5) females. Each sample comprised of pooled *tdRFP*-positive MMECs isolated from multiple mice of the same genotype (see Methods). A. Venn diagrams of the number of significantly altered (differentially expressed) genes (left) and enriched Hallmark Pathways identified by Gene Set Enrichment Analysis (GESA; right) comparing *B-cat+/R1KO* against *B-cat+/WT,* and *B-cat+/DKO* against *B-cat+*/*WT*. The differential expression of a gene was considered significant if it had an absolute fold change greater than 1.5 and an adjusted *P* < 0.05, calculated using Wald test with DESeq2 package. Pathways’ significance determined based on a false discovery rate q-value threshold of 0.05. Circles represent sets of Differentially Expressed Genes (DEGs)/pathways identified in *B-cat^+^*/*R1KO* and *B-cat^+^*/*DKO*; intersecting region shows DEGs/pathways common to both conditions; non-overlapping areas represent DEGs/pathways unique to each condition. B. Lollipop plot displaying normalised enrichment score (NES) from GSEA of the Hallmark Gene Sets comparing *B-cat^+^*/*R1KO* and *B-cat^+^*/*DKO* to *B-cat^+^/WT*. Each lollipop represents a gene set, with NES on the y-axis and the gene sets on the x-axis. Only gene sets that were significant (i.e. FDR q-value of 0.05 or less) in at least one of the conditions where used, with the position of the lollipop indicating the NES, and the size of the bubble corresponding to the -log10 of the FDR-adjusted q-value. Gene ranking for the analysis was based on the signal-to-noise ratio. These plots highlight the most significantly enriched gene sets in each condition, with larger bubbles indicating stronger statistical significance.

Given that combined loss of *Runx1* and *Runx2* resulted in a dramatically accelerated oncogenic phenotype, we were interested in those gene sets that distinguished the *B-cat+*/*DKO* group from the *B-cat+*/*R1KO* one. These fell into two groups. The first refers to gene sets that showed an enriched score when the *B-cat+*/*DKO* dataset was compared against the *B- cat+*/*R1KO* one (Fig. 5B), including MYC, mTORC1, TGFb, EMT, IL6/JAK/STAT3, IL2/STAT5 and WNT/b-catenin signalling (Fig. 5B), possibly contributing to the more aggressive phenotype. The second group includes gene sets negatively enriched in *B-cat+/R1KO* MMECs compared to the *B-cat+/WT* ones yet positively enriched in *B-cat+/DKO* cells. Changes in these pathways (e.g. TNFA, hypoxia, PI3K/AKT/MTOR, ROS and apoptosis) might be key to explaining the dramatic differences observed upon simultaneous *Runx1* and *Runx2* loss (Fig. 5B).

### Deletion of *Runx* in mammary epithelial cells alters the immune microenvironment

To investigate the wider effect of *Runx* deletion on the tumour microenvironment, RNA-seq was also carried out on whole mammary tissue from 9-weeks-old mice. PCA showed that samples within each genotype grouped together (Fig. 6A). In keeping with the *B-cat+/R2KO* cohort having a comparable tumour phenotype to the *B-cat+/WT* (Fig. 2), these two groups were transcriptionally aligned, while the *B-cat+/DKO* one was the most divergent (Fig. 6A). Surprisingly, MetaCore pathway analysis revealed that 7/10 of the most significantly enriched pathways in the *B-cat+/DKO* cohort related to immune pathways (Fig. 6B, lighter bars). These data suggest that loss of *Runx1* and *Runx2* in *Wnt/β-catenin* positive MMECs can profoundly change the immune landscape of surrounding tissues. Deconvolution analysis using two different approaches was applied. First, the Mouse Microenvironment Cell Population counter (mMCP-counter) (41) revealed 16 major immune and stromal cell types using gene expression profiles that identify and quantify different cellular populations. *B-cat+*/*DKO* glands showed a markedly increased presence of immune cells within the mammary gland comprised of macrophages, granulocytes and T cells, including CD8 T cells (Fig. 6C). We then followed up with CIBERSORTx (42) and referenced our RNA-seq data against that of Bach *et al.* (43) who used single cell RNA-seq to fractionate different epithelial, stromal and immune cell types within the murine gland. Three different CD8+ T cell (CD8-1, CD8-2 and CD8-3) and two different CD4+ T cell (CD4-1 and CD4-2) subtypes were identified based on gene expression profiles (43). We noted clear alterations of these subsets in *B-cat+*/*DKO* glands, with a decrease in CD8-1, CD8-2, CD4-2 and CTL populations and an increase in CD8-3 and T-reg cells (Fig. 6D). Our data also demonstrated that *B-cat+*/*DKO* glands are enriched for a macrophage subtype termed Mo3, previously identified as tissue resident macrophages and associated with mammary ducts (44). Altogether, these results indicate that loss of *Runx* in *Wnt/β-catenin*-driven MMECs results in both profound tumour cell changes as well as in significant perturbations to non-tumour cells capable of altering the local environment.

**Fig. 6.**
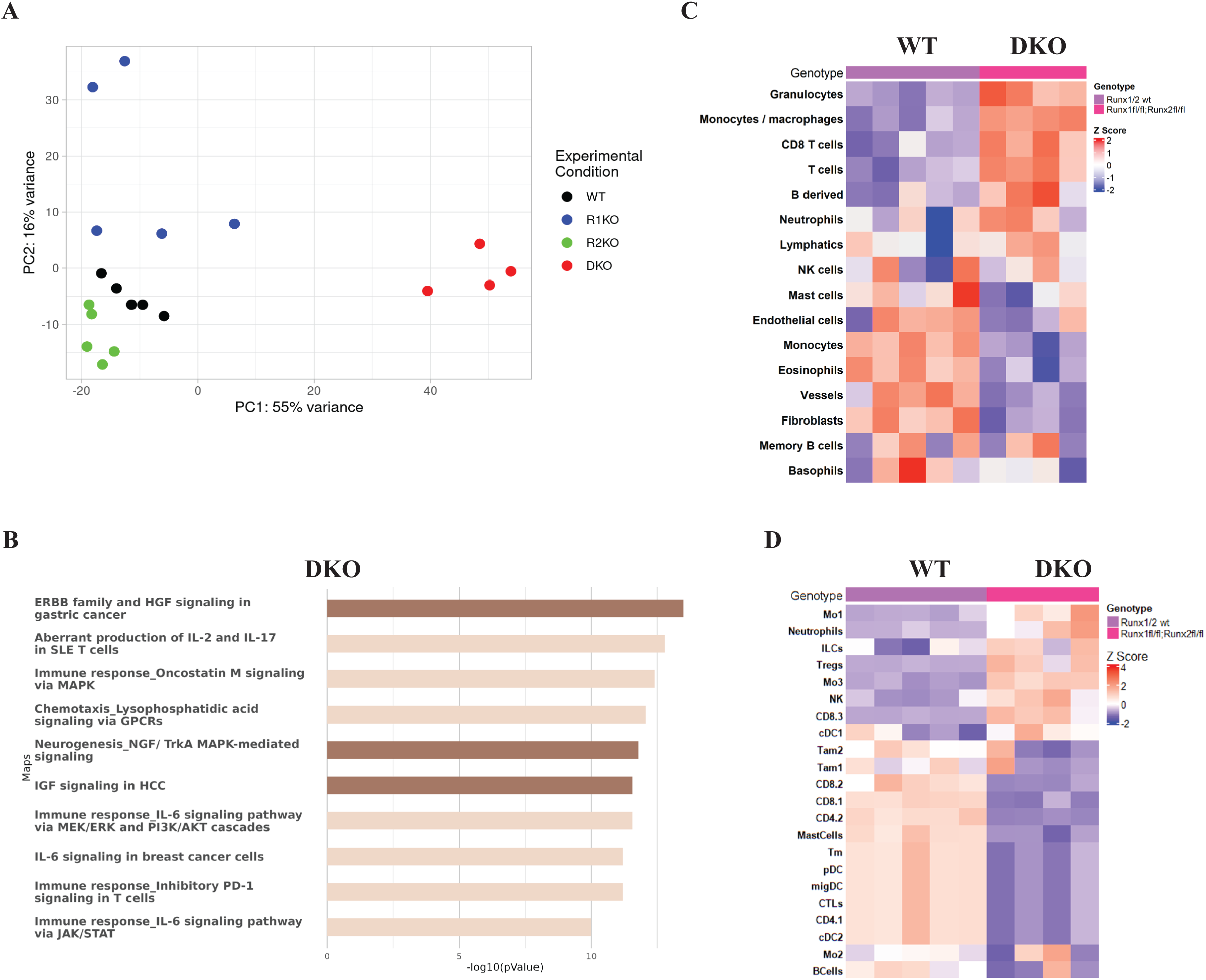
Deletion of *Runx* genes in mammary epithelial cells alters the immune microenvironment. A. PCA plot of the top 500 most variable genes showing RNA-seq expression profiles for whole mouse mammary glands harvested from 9-weeks old *B-cat*+ mice. Each point represents an individual mouse either: *Runx* wildtype (WT; black; *n* = 5), *R1KO* (blue; *n* = 5), *R2KO* (green; *n* = 5), and *DKO* (red; *n* = 4). PC1 and PC2, explaining 55% and 16% of the variance, are on the x- and y-axes, respectively. B. Top enriched pathways from MetaCore’s Canonical Pathway Maps representing signalling and metabolic pathways are shown (for samples as in A). The -log10 of the FDR-corrected p-value indicates the probability of the observed intersection between the DEGs and the pathway gene set occurring by chance, as determined by a hypergeometric test. A higher -log10 p-value suggests a stronger association between the DEGs and the pathway “DKO vs WT” refers to DEGs in the *Runx1*/*Runx2* double knockout compared to wildtype. C. Heatmap displaying z-scored mMCP scores for immune and stromal cell populations in *Runx* wildtype (WT; *n* = 5) versus *Runx1*/*Runx2* double knockout (DKO; *n* = 4) 9- weeks old *B-cat+* mammary glands (as in A). Each column represents an individual mouse. D. Heatmap of CIBERSORTx immune cell fractions in *Runx* wildtype (WT; *n* = 5) versus *Runx1*/*Runx2* double knockout (DKO; *n* = 4) 9-weeks old *B-cat+* mammary glands (as in C and D). CIBERSORTx immune cell fractions calculated using a signature matrix created from single cell sequencing of *BLG-cre;Brca1^f/f^;Trp53^+/-^*murine mammary glands from Bach *et al*. (43). Z-scores are representative of the gene expression relative to the mean expression, indicated by a diverging scale running from blue to red. Each column represents an individual mouse.

## DISCUSSION

*RUNX1* and *CBFB* have been reported to be genetically altered in breast cancer, with many of these changes believed to be loss-of-function mutations (11, 12, 34). In addition, loss of RUNX1 function is associated with disease progression (14, 15, 45). Despite this, it is unclear how the RUNX1/CBFβ complex protects against tumour development. In this study, we demonstrate that *Runx1* loss accelerates mammary tumourigenesis in GEMMs driven by the *PyMT* oncogene or by sustained *Wnt/β-catenin* signalling, providing definitive *in vivo* proof that *Runx1* fulfils a tumour suppressor role in breast cancer, at least in the context of these oncogenic pathways. However, this does not appear to be a universal phenomenon as no tumour acceleration was observed when *Runx1* was deficient in mammary tumours arising from *Trp53* loss. This was surprising given that *Runx1* loss in MMECs had previously been linked with enrichment of genes associated with p53 activation (34). With the observation that arrested development of luminal epithelial cells lacking *Runx1* could be rescued in the absence of p53 resulting in hyperproliferation of ER+ cells, this led to speculate that these events may pre-dispose to breast cancer (34, 46). However, our results suggest that the tumour suppressive functions of *Runx1* and *Trp53* might overlap, consistent with *TP53* and *RUNX1* mutations not co-occurring in METABRIC or TCGA breast cancer datasets. Similarly, others reported that *CBFB* and *TP53* mutations were also mutually exclusive in these datasets (47).

Heightened WNT signalling is a common finding in many breast tumours (48), with aberrant β-catenin activation reported to be preferentially associated with poor clinical outcome in patients with invasive breast cancer, regardless of molecular subtypes (24). Moreover, deregulated WNT signalling is linked with aggressive tumour features, including metastasis and the maintenance of cancer stem cells (49). In the presence of activated *β-catenin*, *Runx1* deficiency promotes tumourigenesis and enhances a stem-like phenotype in both MMECs *in vivo,* as well as HC11 cells *in vitro*. These results agree with others who have shown that *Runx1* drives the differentiation of stem/progenitor mammary epithelial cells (34, 50) and predicted that *RUNX1* loss would result in the expansion of such cells (50).

Although loss of *Runx2* did not change the rate of tumour formation, accelerated tumour onset was observed when *β-catenin* expressing MMECs were deficient for both *Runx1* and *Runx2*. These results indicate that, in the absence of *Runx1, Runx2* may curtail tumour development. Although different family members have unique roles during lineage programming and cell fate decisions, this largely arises from their distinct spatial and temporal expression patterns. Indeed, that the *Runx* genes have significant functional redundancy is not surprising, given they share a common binding site and regulate a common gene set, as established in a number of different systems (51–53).

Our results show that loss of *Runx2* can promote *Wnt/β-catenin*-driven mammary tumourigenesis, but only upon *Runx1* loss. It is possible that, when both genes are functional, *Runx2* represses the effects of *Runx1* by keeping cells in a stem-like state. Accordingly, MMECs expressing activated *β-catenin* and lacking both *Runx1* and *Runx2* are more strongly enriched for gene sets representing mammary stem cells than those with only *Runx1* loss, which themselves show a stronger stem cell programme than control *β-catenin* expressing MMECs. Our finding that combined loss of both genes further enhances this effect suggests a cryptic role for *Runx2* in negatively regulating the stem cell phenotype, only revealed in the absence of *Runx1*. Together, these suggest that both *Runx1* and *Runx2* act in concert to restrain the oncogenic potential and stem-like properties of aberrant *Wnt/β-catenin*.

RNA-seq analysis revealed that upon *β-catenin* activation, many more genes are differentially expressed and many more oncogenic pathways activated when both *Runx* genes are lost. Particularly intriguing are those gene sets negatively enriched in cells that have lost *Runx1* compared to activated *β-catenin* alone, but that are highly enriched when both *Runx* genes are lost (e.g. TNFA and hypoxia). Thus, the rapid tumour onset of multiple clones might be driven both by the expansion of a stem-like population of mammary cells and activation of different oncogenic pathways that are only unleashed in the absence of RUNX function.

An additional finding was the marked enrichment of immune pathways observed in *B- cat+*/*DKO* glands before the appearance of gross lesions. Intriguingly, T-regs were one of the cell types predicted to be enriched in glands with simultaneous epithelial loss of *Runx1* and *Runx2*, suggesting the establishment of an immune suppressive environment. Whilst the mMCP analysis revealed a general T cells increase, refined CIBERSORTx deconvolution indicated that certain T cells subtypes were reduced, whilst others expanded. Future work will understand whether this change in T cell subtypes restricts or promotes tumorigenesis. It is possible that the increase in Mo3 cells displayed by *B-cat+*/*DKO* mice may be secondary to ductal proliferation as others have shown that hormone induced branching can drive expansion of this population (44). Intriguingly, ductal macrophages have been associated with an immunosuppressive function within the gland, characterized by an expression profile aligning with tumour associated macrophages (44, 54). Thus, the altered immune populations might have the potential to influence the cellular milieu that nurtures emerging *B-cat+*/*DKO* tumours, a concept worth further investigation considering that WNT-driven tumours are characteristically renowned to be immune-excluded (55, 56).

In summary, loss of RUNX function within *Wnt/β-catenin*-driven mammary epithelial cells alters the threshold for oncogenic transformation and changes the immune response towards this expanding population of abnormal cells. How these different effects interact and contribute to rapid tumour development observed will be the focus of future studies.

## MATERIALS AND METHODS

### Animal procedures

Mouse experiments were performed with ethical approval from the University of Glasgow Animal Welfare and Ethical Review Board under the Animal (Scientific Procedures) Act. Mice were housed in a pathogen-free facility in individual ventilated cages on a 12h light/dark cycle with continual access to food, water and environmental enrichment. Genotyping was carried out by Transnetyx (Cordova, US). Mouse euthanasia was humanely performed.

### Transgenic lines and genetic models of mammary cancer

Mouse Mammary Tumour Virus (*MMTV*)*-Cre* (57) and *MMTV*-Polyomavirus middle T antigen (*PyMT*) (26) lines were provided by WJ Muller. Tandem dimer red fluorescent (*Gt(ROSA)26Sortm1Hjf*) knock-in *Cre*-reporter line (hereafter *tdRFP*) was obtained from the European Mouse Mutant Archive (28). Beta-lactoglobulin (*BLG*)*-Cre* (30); *Ctnnb1^wt/lox(ex3)^* (29); *Trp53^fl/fl^* (58); *Runx1^fl/fl^* (59) and *Runx2^fl/fl^* (18) alleles as previously published. *MMTV-PyMT* mice were crossed with *MMTV-Cre;Runx1^fl/fl^*to generate *PyMT+/R1KO* and *PyMT+/WT* lines, backcrossed 10 generations to FVB/N [Charles River, UK]. *BLG-Cre;Ctnnb1^wt/lox(ex3)^* mice, bred with *Runx1^fl/fl^* and *Runx2^fl/fl^* lines, were intercrossed to establish cohorts maintained on an outbred background (C57BL/6; FVB/NCrl) as confirmed by Transnetyx MiniMUGA (60). *BLG-Cre;Trp53^fl/fl^*mice, bred with *Runx1^fl/fl^* and *Runx1^wt/wt^* colonies, were backcrossed 10 generations to FVB/N [Charles River, UK]. To maximize *BLG-Cre* expression (30), females were mated at 11-13 weeks for two rounds of pregnancy, except for three *BLG-Cre;Ctnnb1^wt/lox(ex3)^;Runx2^fl/fl^*(*B-cat+/R2KO*) females mated at 7-9 weeks.

### Tumour onset, tumour volume and clinical endpoint analyses

Mice were palpated for tumour formation twice weekly. Tumour onset was defined as lesions ≥2mm and ≤7mm. Mice with tumours >7mm at time of notice were excluded from analysis. Tumours were measured with calipers and volume (mm^3^) calculated as follows: Volume = (width x width x length)/2. Mice were euthanised when tumours reached endpoint (largest diameter at 15mm or ulcerated) and number of tumour-bearing glands (of 10 in total) counted.

### Wholemounts and histology

Virgin females were sacrificed at 6- and 9-weeks. For wholemount staining, right-side abdominal (#4R) mammary glands were excised, spread on glass slides, air-dried and placed overnight in Carnoy’s solution (6-parts EtOH/3-parts CHCl_3_/1-part glacial acetic acid). Slides were washed in EtOH/15 min (70%, 50%, 25%), rinsed in dH20/10 min and stained overnight in Carmine Alum. Glands were washed in EtOH/15 min (70%, 95%, 100%), cleared in Xylene, mounted with Pertex Mounting Media and coverslipped. Wholemounts were photographed with a Zeiss stereomicroscope. For H&E staining, left abdominal glands (#4L) were formalin-fixed, paraffin-embedded, sectioned at 4µm and mounted on glass slides by the CRUK Scotland Institute Histology Department.

### Mouse mammary epithelial cell (MMEC) extraction

Mammary glands (with lymph nodes removed) were dissected, placed in serum-free working solution - 1X F-12 Nutrient Mixture (Ham), 1X Penicillin-Streptomycin-Glutamine (PSG) - chopped [Mcllwain] and tissue paste incubated (1.5 h/37°C) with serum-free digestion solution - 1X Ham, 1X PSG, 10% Collagenase (300U/ml)/Hyaluronidase (100U/ml) - in a shaking incubator (100rpm). Organoids were collected through centrifugation (350g/5 min), while red blood cells lysed in 0.8% NH_4_Cl (RT/5 min) and removed by aspiration. To remove stromal cells, pellet was re-suspended in 10% Foetal Bovine Serum (FBS)-supplemented working solution, plated and incubated (37°C/1 h). Floating epithelial organoids were isolated and centrifuged (350g/5 min). Pellet was washed with 1X Phosphate-Buffered Saline (PBS), re- suspended in 0.5-1ml dissociation solution - 1X PBS, 0.25% Trypsin, 1mM EGTA, 0.1mg/ml DNase I recombinant, RNase-free 10000U - and incubated (37°C/10 min) to obtain a single cell suspension. MMECs were purified by filtering (40µm) and centrifugation (350g/5 min).

### Flow cytometric analysis and fluorescence-activated cell sorting (FACS)

MMECs were isolated from *PyMT*+/*WT* and *PyMT+/R1KO* mice at tumour notice (average of 61 and 50 days, respectively). An equivalent time-point (∼60 days) was chosen for *PyMT-* negative cohorts. One abdominal gland (#4R) was retained for histology. Remaining glands were analysed as individual samples (except for thoracic glands combined due to their overlap, i.e. #2/3R and #2/3L). 7 samples were analysed from *PyMT+* cohorts (#1R, #1L, #2/3R, #2/3L, #4L, #5R and #5L), 5 samples from *PyMT-*negative mice due to scarcity of MMECs in #1 and #5 glands, which were combined (i.e. #1RL, #2/3R, #2/3L, #4L, #5RL). Data were run on an Attune Nxt Flow Cytometer [Invitrogen] and analysed with FlowJo (version 9.9.6). For FACS of *tdRFP*-positive MMECs from *B-cat+* cohorts, ten glands were pooled from each 9-week mouse, MMECs isolated, re-suspended in FACS buffer - 1X PBS, 2mM EDTA pH 8, 1% FBS - and stained with DAPI/5 min. For each experiment, *tdRFP*-negative MMECs were used to set gates. Cells were collected using a BD FACSAria™ III Cell Sorter.

### Colony forming cell assay

MMECs from 7-9-weeks *B-cat+* mice were plated onto 24-well plates (5×10^3^ viable cells/well), embedded in 25µl of Matrigel and cultured with 500µl of medium - 1X EpiCult-B Basal Medium Mouse, 10% EpiCult-B Proliferation Supplements, 5% FBS, 1% PSG, 4µg/ml Heparin, 10ng/ml mouse epidermal growth factor (mEGF), 10ng/ml mouse fibroblast growth factor-2 (mFGF-2). MMECs were extracted from all ten glands per mouse and fed with 500µl of fresh medium at day 3. Representative pictures were taken at day 7-9 and number of acinar- and solid-like colonies counted using a Olympus CKK41 microscope (4x lens).

### HC11 cell assays

HC11 and HEK293T cells [ATCC] were maintained in RPMI 1640 (+10% FBS, 1% PSG, 5µg/ml Insulin, 10ng/ml mEGF) or DMEM (+10% FBS, 1% PSG) media, respectively. For 2D growth assays, trypsinised cells were plated (5×10^3^ viable cells/ml) and supplemented with fresh medium every 2-3 days. For each condition, four 24-wells were counted every 24h for 2-4 experimental repeats. For mammosphere assays, 5×10^3^ viable HC11 cells/ml in mammosphere medium - RPMI 1640, 1% PSG, 20ng/ml mEGF, mFGF-2, 4µg/ml heparin, 1X B27 Supplement - were cultured in 24-well Ultra-Low Attachment plates at 5% CO_2_/37°C. Cells were supplemented with mEGF and mFGF-2 (20ng/ml) every 3 days. Recombinant mouse WNT3A protein [100ng/ml, Abcam] was added to mammosphere medium and supplemented every 3 days. Number of mammospheres per well was counted at day 7 using an Olympus CKX41 Inverted Phase Contrast Microscope (10x lens).

### *Runx1* overexpression & CRISPR/CAS9 gene targeting

For *Runx1* overexpression, HEK293T cells were transfected with 10µg of pBABE-Puro (Empty or *mRunx1P1*), 7.5µg of psPAX2 [Addgene] and 4µg of pVSV-G [Addgene] plasmids using calcium phosphate with medium replacement 24h post-transfection. Virus-containing media, filtered through 0.45µM, were used to transduce HC11 cells with 10µg/ml polybrene. Cells were selected with 1.5µg/ml Puromycin. For CRISPR/CAS9-mediated targeting, guide RNAs (gRNAs) targeting *Runx1* and *Runx2* were designed using the Zhang Lab tool [MIT, Boston, MA]. pX-459-Puro (Empty or *Runx1*-targeting gRNAs) constructs were used to transfect HC11 cells using Lipofectamine™ 2000 Transfection Reagent. Three different *Runx1*-targeting gRNAs were tested (*mhRunx1*, *mRunx1_A*, *mRunx1_B*). Three independent *Runx1*-deleted (R1KO_a, R1KO_b, R1KO_c) HC11 clones were generated using the *mRunx1_B* gRNA sequence (5’-CGGTGCGCACTAGCTCGCCA). For *Runx1/Runx2* deletion, HC11 cells were transfected with pX-459-Puro (Empty or *Runx2*-targeting gRNA, i.e. *mhRunx2*: 5’-GAGCGACGTGAGCCCGGTGG) constructs, then transduced with lentiCRISPRv2-Neo (Empty or *Runx1*-targeting gRNA, i.e. *mRunx1_B*) vectors as above. Upon antibiotic selection (800µg/ml G418 Sulphate/Geneticin) and single-cell cloning, an independent *Runx1*-deleted (R1KO_d), two independent *Runx2*-deleted (R2KO_a, R2KO_b) and two independent *Runx1/Runx2*-deleted (DKO_a, DKO_b) HC11 clones were generated.

### Immunoblotting and antibodies

HC11 cells were lysed in Pierce RIPA Buffer. Protein extracts, resolved on 4-12% NuPAGE Novex Bis-Tris 1.0mm Mini Protein Gels [Thermo Fisher Scientific], were transferred to Hybond-ECL nitrocellulose membranes [Amersham]. The following antibodies were used: RUNX1 [#8529, D4A6, Cell Signaling; 1:1000]; RUNX2 [#8486, D1H7, Cell Signaling; 1:1000]; GAPDH [#3683, 14C10, Cell Signaling; 1:1000] and horseradish peroxidase-conjugated anti-rabbit secondary antibody [#7074, Cell Signaling; 1:10 000]. Densitometry analysis performed using Image Lab software (Fiji ImageJ software version 1.53).

### RNA sequencing

RNA extraction was performed using the RNAqueous™-Micro Total RNA Isolation Kit [Invitrogen] and residual genomic DNA removed using the DNA-free™ system*. tdRFP-* positive sorted cells were pooled from 3-6 individual mice and combined into a single 20µl RNA sample. NanoDrop 2000 Spectrophotometer was used to quantify and assess purity of extracted RNA. RNA quality was assessed on an Agilent 2200 TapeStation System using RNA ScreenTape Analysis. Samples with RNA integrity number (RIN) values ≥ 7 were used wherever possible (18/19 samples in Fig. 6) and RIN values ≥ 6 (8/14 samples) for *tdRFP+* FACS cells (Fig. 3 and 5). RNA-seq libraries were prepared using SMARTer Stranded Total RNA-Seq Kit v2 - Pico Input Mammalian [Takara Bio]. Library quality was assessed using Agilent 2200 TapeStation System with a High Sensitivity D1000 ScreenTape Analysis and quantity measured with a Qubit Fluorometer. Libraries were sequenced by W. Clarke at CRUK Scotland Institute Molecular Technologies Facility via the Illumina NextSeq 500 System with paired-end (2×36) sequencing using the MiniSeq High Output Reagent Kit (75-cycles). Illumina sequence data were demultiplexed using bcl2fastq (v2.20). Quality checks on raw RNA-Seq data files were done using FastQC (https://www.bioinformatics.babraham.ac.uk/projects/fastqc/) and FastQ Screen (61). RNA-Seq paired-end reads were aligned to the GRCm38.93 version of the mouse genome and annotation (62) using HiSat2 (63) and sorted using Samtools (64). Aligned genes were identified using HTSeq (65). Expression levels were determined and statistically analysed using the R environment (https://www.r-project.org/) and utilising packages from the Bioconductor data analysis suite (66). Differential gene expression was analysed based on the negative binomial distribution using DESeq2 (67) and adaptive shrinkage using Apeglm (68). Identification of enriched biological functions was achieved using g:Profiler (69), GESA from the Broad Institute (70) and MetaCore from Clarivate Analytics (https://portal.genego.com/). Computational analysis was documented at each stage using MultiQC (71), Jupyter Notebooks (https://eprints.soton.ac.uk/403913/) and R Notebooks (https://posit.co/).

### CIBERSORTx Analysis

Signature matrix was generated from an annotated single-cell RNA-seq dataset of murine glands (43) to provide a reference to impute cell fractions from gene expression profiles. Signature matrix and normalised bulk RNA-seq datasets were run through CIBERSORTx to deconvolute the bulk RNA-seq datasets (https://cibersortx.stanford.edu). CIBERSORTx job was run under Impute Cell Fractions settings, relative run mode, >100 permutations for significance analysis, and with quantile normalisation disabled. CIBERSORTx results were exported as CSV files and imported into R Studio (v 4.3.2) for visualization.

### mMCPCounter Analysis

mMCPCounter (mMCPcounter.estimate) is a function within the mMCPcounter (v 1.1.0) R package in R studio (41). mMCPcounter.estimate was used to generate scores for 16 cell populations. Heatmaps of cell population Z-scores were plotted using the ComplexHeatmap (v 2.14.0) R package. To compare statistics of the boxplots for the different cell populations, the stats (v 4.2.2) R package was used to perform a t test between each of the conditions.

### Statistical analysis

Statistical analyses performed using GraphPad Prism (version 10). Specific statistical tests used are indicated in figure legends. All error bars represent mean ± SD, unless otherwise indicated.

## DATA AVAILABILITY

All data generated and/or analysed during the current study are available from the corresponding authors. Full RNA-seq datasets will be publicly available on the Sequence Read Archive database when accepted for publication.

## ACKNOWLEDGMENTS

Authors wish to acknowledge all funding sources listed below. We thank the Advanced Technologies and Core Services at the CRUK Scotland Institute, especially the Biological Services Unit, the Histology Facility, William Clarke and Molecular Technologies, the Flow Cytometry Facility, and the Beatson Advanced Imaging Resource (BAIR). We are grateful to all past and present members of the Blyth lab for useful discussion and continued support during the project. Thank you to Barry Gusterson, Joanna Morris (University of Glasgow) and Matthew Smalley (Cardiff University) for advice and discussion, and to Catherine Winchester (CRUK Scotland Institute) for critical review of the manuscript and advice on research integrity. Schematics were prepared using BioRender with licence to K Blyth.

## AUTHOR CONTRIBUTIONS

*Conceptualization*: AIR, KJC, ERC, KB

*Methodology*: AIR, KS, NF, KG, CB, HH, CN, KB

*Investigation*: AIR, KS, NF, DA, KJC

*Formal Analysis*: AIR, KS, RS, AL, KG, CB, ALY, HH, PDA, EWR, CJM, PDD, ERC, KB

*Data Curation*: RS, AL

*Visualization*: AIR, KS, RS, AK, AL, ERC

*Validation*: AK, AL, FG, KJC

*Resources*: ALY, PDD

*Supervision*: PDA, EWR, CJM, PDD, KJC, ERC, KB

*Funding Acquisition*: KB, EWR, CJM, PDD

*Writing – original draft*: AIR, ERC, KB

*Writing – reviewing/editing*: AIR, RS, AK, AL, KJC, ERC, KB

All authors have approved the final version of the manuscript.

