## Supplemental data for "*Runx1* and *Runx2* act in concert to suppress *Wnt/β-catenin*-driven mammary tumourigenesis"

### Supplementary Fig. 1

A

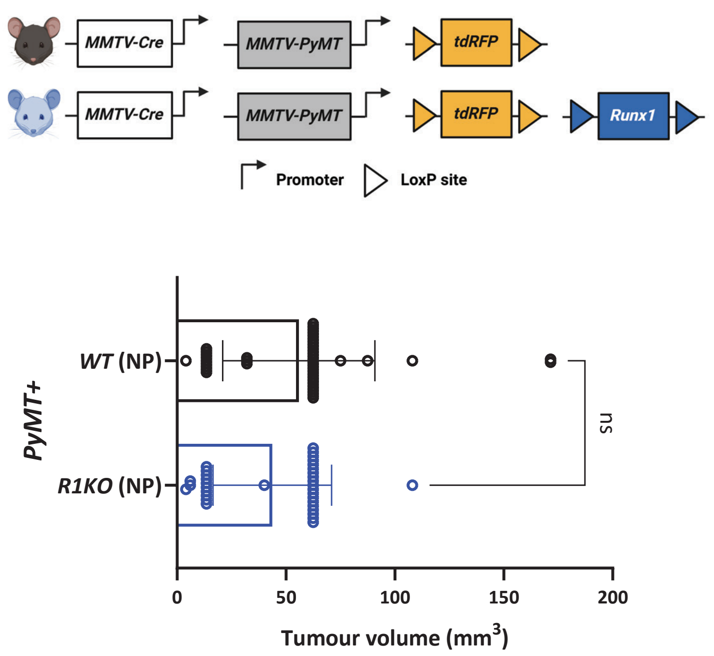


B

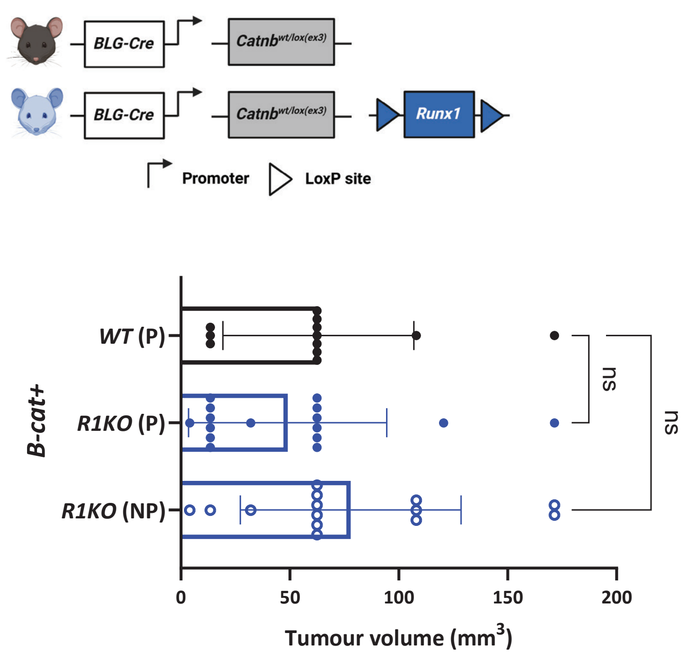


C

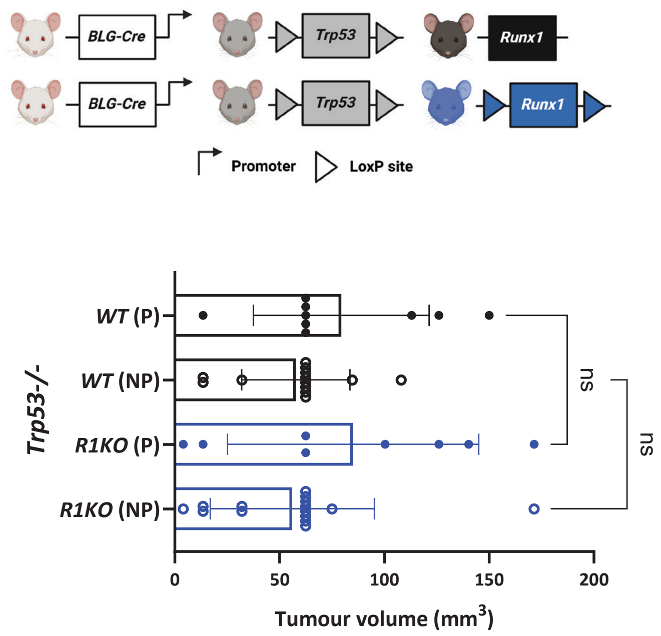


D

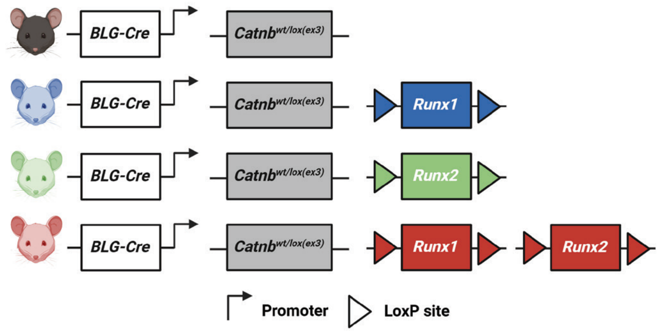

**Supplementary Fig. 2 Loss of *Runx1* and *Runx2* perturbs mammary gland development in the presence of activated Wnt/β-catenin signalling.**

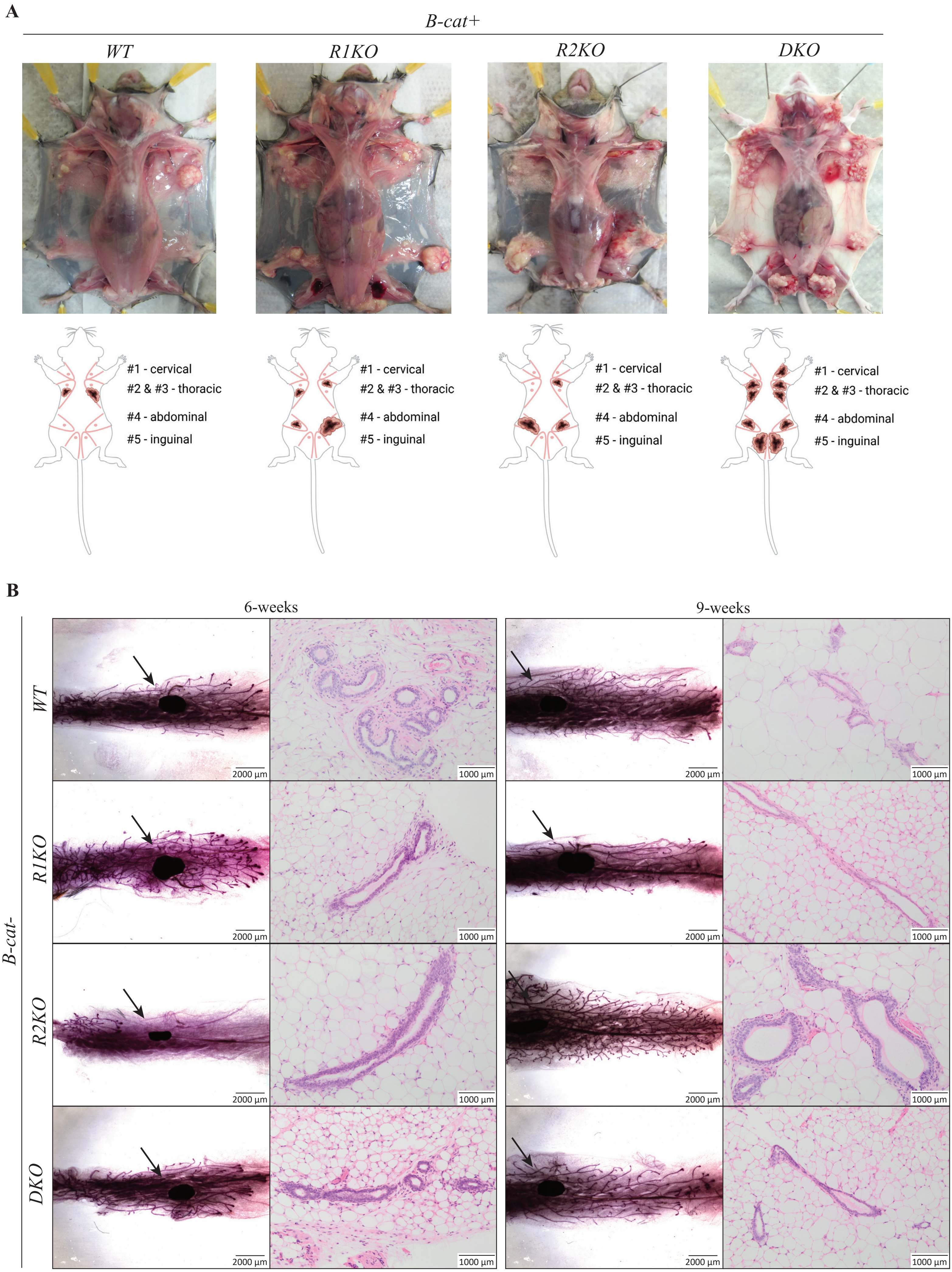

**Supplementary Fig. 3 Altered *Runx1* expression does not affect the phenotype of HC11 cells in 2D cultures.**

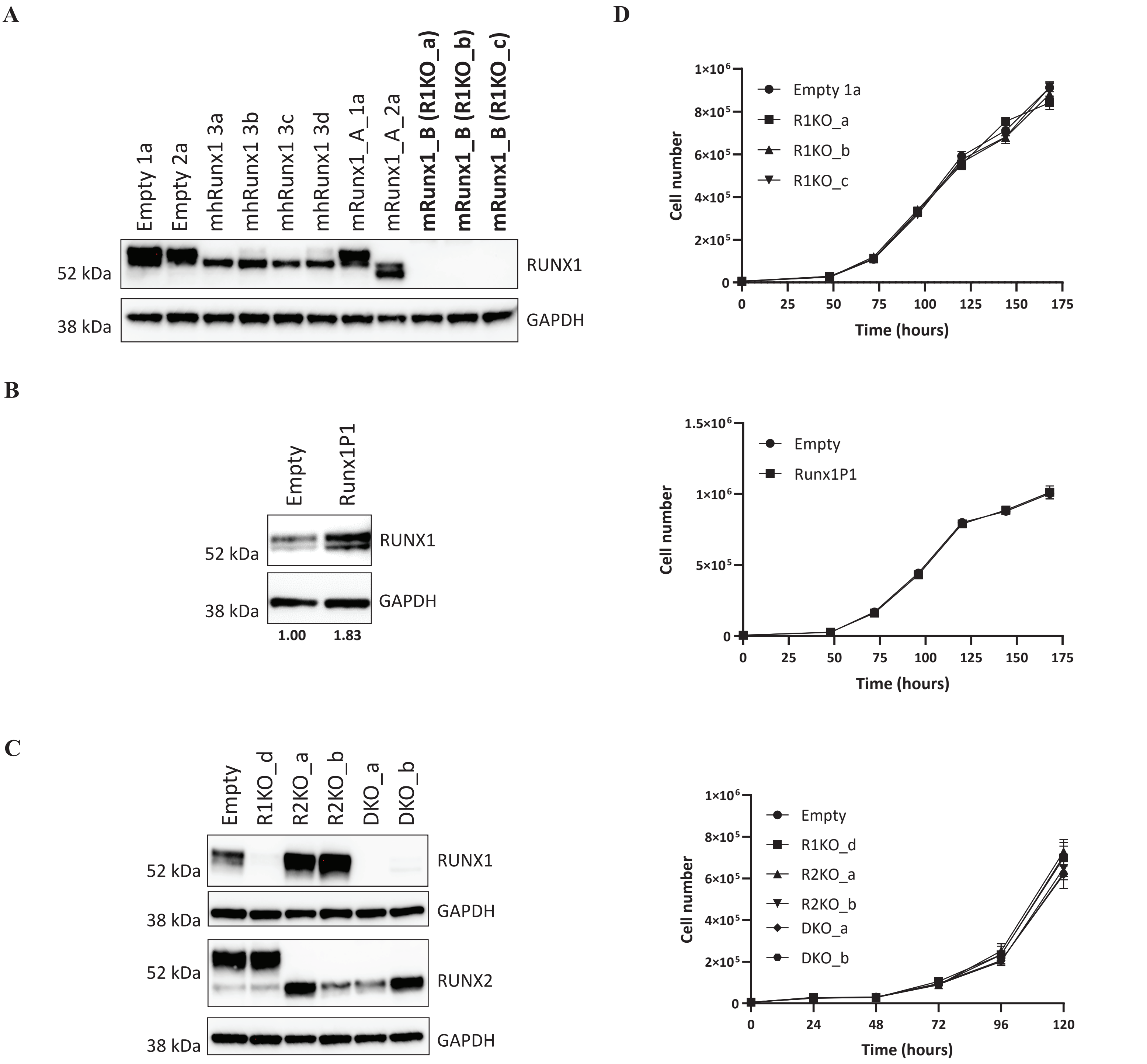

### SUPPLEMENTARY FIGURE LEGENDS

#### Supplementary Fig. 1.

A. Schematic of mouse cohorts (top) and tumour volume (bottom) of *PyMT*+ cohorts as in Fig. 1A ($n = 44$; *WT*; $n = 34$; *R1KO*) at date of tumour notice. Average tumour volume with standard deviation shown: 56 $mm^3$ (± 34.9) for *WT* and 44 $mm^3$ (± 27.2) for *R1KO*. NP, nulliparous. Statistical analysis performed with two-tailed unpaired t-test; ns, non-significant ($P > 0.05$).

B. Schematic of mouse cohorts (top) and tumour volume (bottom) of *B-cat*+ cohorts as in Fig. 1C ($n = 12$, *WT*-parous; $n = 16$, *R1KO*-parous; $n = 14$, *R1KO*-nulliparous) at date of tumour notice. Average tumour volume with standard deviation shown: 63$mm^3$ (± 43.8) for *WT*-parous, 49$mm^3$ (± 45.5) for *R1KO*-parous and 78$mm^3$ (± 50.7) for *R1KO*-nulliparous. Statistical analysis performed with ordinary one-way ANOVA test with Dunnett's multiple comparisons test; ns, non-significant ($P > 0.05$).

C. Schematic of mouse cohorts (top) and tumour volume (bottom) of *BLG-Cre;Trp53*$^{fl/fl}$ cohorts as in Fig. 1D ($n = 9$, *WT*-P; $n = 13$, *WT*-NP; $n = 8$, *R1KO*-P; $n = 15$, *R1KO*-NP) at dates of tumour notice. Average tumour volume with standard deviation shown. Parous (P) and nulliparous (NP) *WT* mice: 79$mm^3$ (± 41.9) and 58$mm^3$ (± 25.8), respectively. *R1KO*-P and *R1KO*-NP mice: 85$mm^3$ (± 59.9) and 56$mm^3$ (± 39.2), respectively. Statistical analysis performed with ordinary one-way ANOVA test with Sidak's multiple comparisons test; ns, non-significant ($P > 0.05$).

D. Schematic of mouse cohorts shown in Fig. 2A.

**Supplementary Fig. 2 (relative to Fig. 2).**

A. One representative picture shown per genotype ($n \geq 9$ mice per genotype) with diagrams below illustrating mammary gland location and necropsy examination of *B-cat+* cohorts at clinical endpoint (as in Fig. 2B). Note the distinct formation of multiple, sometimes coalescing tumours affecting almost all mammary glands in nulliparous *B-cat+/DKO* mice compared to other genotypes (as taken from parous cohorts).

B. Wholemount and H&E images of abdominal mammary glands from 6- and 9-weeks old *BLG-Cre;Runx1*$^{wt/wt}$ (*WT*; $n$ = 5 for 6-weeks; $n$ = 6 for 9-weeks), *Runx1*$^{fl/fl}$ (*R1KO*; $n$ = 7 for 6-weeks; $n$ = 7 for 9-weeks), *Runx2*$^{fl/fl}$ (*R2KO*; $n$ = 4 for 6-weeks; $n$ = 5 for 9-weeks) and *Runx1*$^{fl/fl}$*;Runx2*$^{fl/fl}$ (*DKO*; $n$ = 7 for 6-weeks; $n$ = 8 for 9-weeks) cohorts (all without *Ctnnb1*$^{wt/lox(ex3)}$). One representative wholemount and H&E shown per genotype. Scale bars of wholemounts = 2mm; H&E images = 1mm. Arrow depicts lymph node.

**Supplementary Fig. 3 (relative to Fig. 4).**

A. Western blot of HC11 cells transfected with empty vector (Empty1a, Empty2a) or carrying three different *Runx1*-targeting guide RNA (gRNA) sequences (*mhRunx1*, *mRunx1*_A and *mRunx1*_B). RUNX1 depletion achieved using the *mRunx1*_B (depicted in bold) in three independent clones (R1KO_a, R1KO_b, R1KO_c) used in 2D (Supplement Fig. 3D) and mammosphere assays (Fig. 4A). GAPDH used as loading control.

B. Western blot of HC11 cells transduced with pBABE-Puro *mRunx1P1* or pBABE-Puro empty vector. Densitometry analysis revealed an almost 2-fold increase in RUNX1 protein levels in *RunxP1* cells (values below). GAPDH used as loading control. Relative to Fig. 4B and Supplement Fig. 3D.

C. Western blots of HC11 cells CRISPR-deleted for *Runx1* (R1KO_d), *Runx2* (R2KO_a, R2KO_b) or *Runx1/Runx2* (DKO_a, DKO_b). Note the non-specificity of the lower band in the RUNX2 blot (on-target gene editing was confirmed by genomic DNA sequencing). GAPDH used as loading control. Relative to Fig. 4C and 2D growth shown in Supplement Fig. 3D.

D. 2D growth curves of three independent *Runx1*-deleted HC11 clones (R1KO_a, R1KO_b and R1KO_c; top panel as in Supplement Fig. 3A), one *Runx1*-overexpressing clone (Runx1P1; middle panel, as in Supplement Fig. 3B), and an independent *Runx1*-deleted clone (R1KO_d), two independent *Runx2*-deleted clones (R2KO_a, R2KO_b) and two independent *Runx1*/*Runx2*-deleted clones (DKO_a, DKO_b; bottom panel, as in Supplement Fig. 3C). For each experiment, HC11 clone(s) were compared to their relative empty vector control line (Empty). Results show mean with error bars representing standard deviation and are representative experiments of $n$ = 2-4 experimental repeats per condition.
